# Extraction of a recombinant full-length NADPH-cytochrome P450 oxidoreductase from bacterial membranes: effect of detergents and additives

**DOI:** 10.1101/279216

**Authors:** Sara A. Arafeh, Azamat R. Galiakhmetov, Elizaveta A. Kovrigina, Eric Fellin, Evgenii L. Kovrigin

**Affiliations:** Chemistry Department, Marquette University, P.O. Box 1881, Milwaukee, Wisconsin 53201, United States

**Keywords:** POR, CPR, CYPOR, expression, alkylamine, polyamine

## Abstract

NADPH-cytochrome P450 oxidoreductase (POR) is a membrane protein in the endoplasmic reticulum of eukaryotic cells. POR is as a key reducing partner for a number of cytochrome P450 proteins involved in different metabolic degradation and signaling pathways. Preparation of the full-length recombinant POR expressed in bacteria has been reported and, typically, involved the use of Triton X-100 detergent for extraction of the overexpressed POR from bacterial membranes. However, extraction efficiency is always relatively low hindering structural studies, particularly—the NMR spectroscopy requiring isotopic enrichment. In this paper, we assessed the effect of a variety of detergents and additives on the efficiency of the membrane-extraction step in POR preparation protocol. We evaluated non-ionic detergents with the variable hydrophobicity (Triton X-100, X-114, and X-405) and structure (Triton X-100, TWEEN-20, Brij-35), a zwitterionic/non-ionic detergent combination (Triton X-100 and CHAPS), as well as a range of alkylamines and polyamines as additives to the conventional extraction buffer containing Triton X-100. None of the detergents or detergent-additive combinations yielded better extraction efficiency than the conventional protocol with the Triton X-100. Lack of variation of the extraction yield allows to hypothesize that the conventional protocol extracts all of the available natively-folded monomeric POR while the remaining fraction is possibly an unfolded aggregated POR, which did not insert in the membranes during expression. We propose that the yield of soluble POR may be increased by a careful optimization of expression conditions while monitoring the distribution of POR between soluble and insoluble fractions in the detergent extraction step.

## Introduction

The NADPH-cytochrome P450 oxidoreductase (POR; also known as CYPOR or CPR) is a protein of the endoplasmic reticulum attached to the membrane through a short N-terminal hydrophobic domain.^1^To isolate and purify POR for biochemical and structural investigations, it was extracted from the membrane using proteases or detergents^2^. However, proteolytic cleavage of the membrane domain results in a truncated protein that is unable to interact with its electron acceptors, cytochromes P450.^1,2,3^Therefore, to prepare the full-length POR including its transmembrane region, the protein has to be extracted from the cellular membranes using a detergent such as Triton X-100, Emulgen 911, or cholate followed by multiple steps of column chromatography.^2,4,5,6,7^ The yields of the recombinant full-length POR, however, remain very low hindering structural studies that require significant amounts of a pure recombinant protein. In this paper, we report a comprehensive analysis of one of the yield-limiting steps of the POR preparation protocol—extraction of the membrane-bound recombinant POR from the bacterial membrane preparation.

## Materials and Methods

### Chemicals and reagents

All chemicals and reagents were purchased from commercial sources and used without further purification. Water for solutions and buffers was purified using Branstead Water Mixed Bed Deionizer. The solubilization buffer (KEG buffer), pH 7.2, was prepared to contain 25 mM KH_2_PO_4_, 100 mM NaCl, and 10% glycerol.

### Expression of POR

The ampicillin-resistant pOR263 plasmid containing the His-tagged full-length POR Q157C/Q517C on a cysteine-less background^8^ was transformed into *E.coli* C41 (DE3) competent cells using a heat shock at 42°C for 40 seconds. The transformed cells were allowed to recover in a Terrific Broth (TB) medium in a culture tube with shaking at 220 rpm for 1 hour at 37°C. The 100 uL aliquot of the transformed cell suspension was plated on an agar-TB plate prepared with 100 ug/mL ampicillin and incubated at 37°C overnight. The next day, a single colony was inoculated into 5 ml of TB medium containing 100 ug/mL ampicillin and incubated at 37°C with shaking at 220 rpm for 3 hours. Next, the culture was transferred into a larger volume of sterile TB medium containing 100 ug/mL ampicillin and incubated at 37°C with shaking at 220 rpm overnight. On the following day, the overnight culture was transferred to a larger volume of the TB medium containing TB salts (72 mM KHPO_4_ and 17 mM KH_2_PO_4_) and 100 ug/mL ampicillin. The culture was shaken at 220 rpm and 37°C until the optical density at 600 nm (OD_600_) reached 0.9 au (approx. 3 hours). The cell culture was cooled on ice briefly and was further shaken at room temperature for 10 minutes. A portion of the cell culture was set aside to represent the non-induced cell culture. To induce overexpression of the full-length POR in the rest of the cell culture, IPTG was added to 0.5 mM and cells were further incubated at 18°C with shaking at 220 rpm overnight. In morning of the following day, the OD_600_ of the culture was in a range of 3-6 au. The cells were harvested by centrifugation at 5,000 g, 4°C for 15 minutes. The cell pellet was resuspended in KEG buffer (25 ml per 1 L of original cell culture) containing 1 mM PMSF and 1 mM EDTA.

### Preparation of membrane samples

To initiate cell lysis, lysozyme was added to 25 µg/mL to the cell pellet suspension. The lysis was allowed to proceed for 1 hour at 4°C followed by sonication on ice (Branson 450 Sonifier; 50% duty cycle, power setting of 8). The samples enriched in POR bound to microsomal membranes were prepared by a series of centrifugation steps resulting in the pellet 2 preparation (Figure 1). Pellet 2 was resuspended in KEG buffer with 1mM PMSF and 1mM Tris(2-carboxyethyl)phosphine (TCEP), aliquoted, flash-frozen, and stored at -80°C.

**Figure 1.**
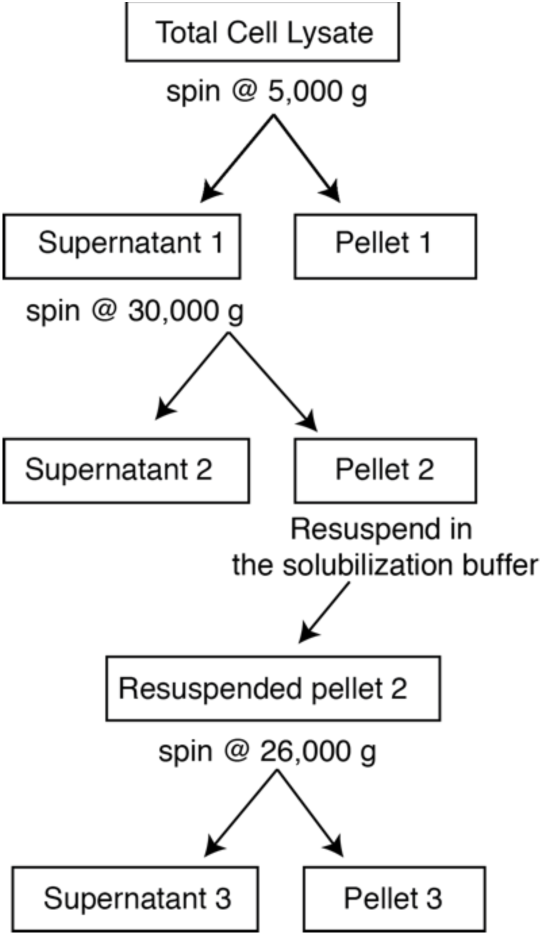
A workflow for isolation of the membrane-bound full-length POR. The first centrifugation sediments cell walls and genomic DNA. Membrane proteins, including POR, remain soluble, associated with microsomes, lipid vesicles formed from bacterial lipid membranes upon sonication. Next, supernatant 1 is ultra-centrifuged at 30,000 g pelleting the membrane vesicles forming pellet 2 (soluble bacterial proteins remain in supernatant 2). Resuspension of pellet 2 in the solubilization buffer (containing various detergents) is followed by another ultra-centrifugation at 26,000 g to separate the solubilized full-length POR (supernatant 3) from POR that remains associated with membrane vesicles or irreversibly aggregated (pellet 3)

### Extraction of POR from membrane samples

All experiments with extraction of POR by different detergents used identical aliquots of the pellet 2 preparation. In a typical extraction experiment, an aliquot of pellet 2 was thawed and sonicated briefly to achieve complete resuspension of the pellet. A specific detergent with/without an additive was added and incubated overnight at 4°C while stirring. The extent of solubilization of full-length POR was assessed by the final centrifugation step (Figure 1) to obtain the resulting supernatant 3 (contained the solubilized POR) and pellet 3 (contained insoluble POR). In the extraction trials when detergents were tested individually, they were added to 2% (v/v) in the solubilization buffer. When detergents were mixed, they were added to 0.5% (v/v) in the buffer. Alkylamine and polyamine additives were used at a final concentration of 50 mM following Yasui et al.^9^

## Results and Discussion

The goal of this work was to increase extraction efficiency of the recombinant POR from bacterial membranes. Conventional protocols for extraction of full-length POR used Triton X-100 but the overall efficiency did not exceed approx. 10%^4,5^. To find ways to increase the protein yield, we varied properties of the solubilization buffer (Figure 1) by using different detergents and their combinations. Because our aim was to enable NMR studies that will utilize labeling of surface-exposed cysteines in POR, we used the full-length POR construct that was engineered to remove all endogenous cysteines and contained two surface-exposed cysteine residues at positions 157 and 517^8^. This construct, the full-length Q157C/Q517C POR with a cysteine-less background, will be referred in the following as “POR".

The membrane-bound POR sample was prepared as described in Materials and Methods from a single batch of the *E. coli* cell culture to ensure that all detergent-extraction trials use the same starting material. Figure 2 demonstrates overexpression of POR in the *E. coli* bacterial cells. The expression level of POR is not particularly high, which is the major motivation for improving efficiency of POR extraction for the preparation of large NMR samples.

**Figure 2.**
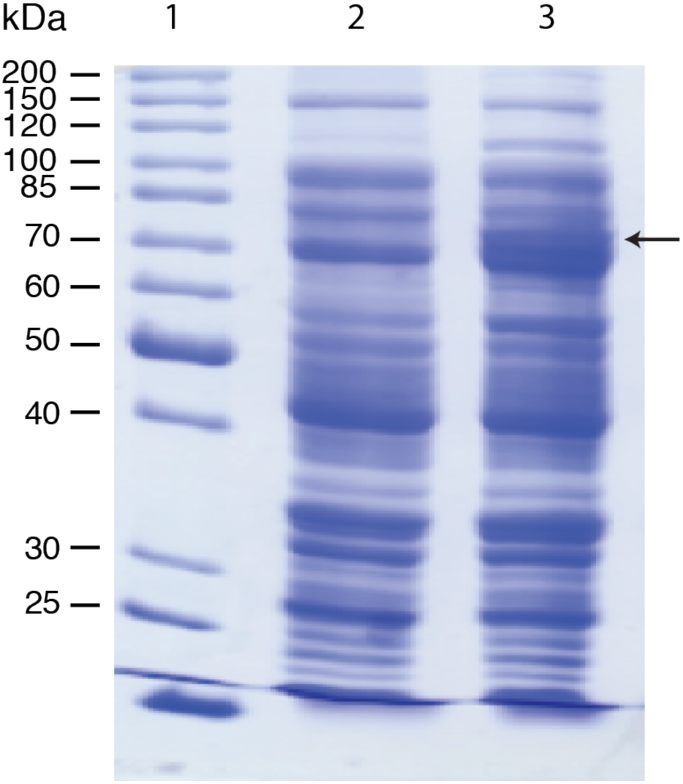
The full-length POR overexpression in a bacterial culture. PageRuler Unstained Protein Ladder, lane 1; the non-induced cell culture, lane 2; the IPTG-induced culture, lane 3. The band of the full-length POR is indicated with an arrow (78kDa).

Since the traditional protocol for POR extraction uses Triton X-100 in the solubilization buffer, the Triton X-100 extraction was performed as a reference experiment^5^. Figure 3. A demonstrates the (typical) low efficiency of extraction of membrane-bound POR from pellet 2. To ensure that POR was not degrading during overnight incubation with detergents, the sample of pellet 2 resuspended in the solubilization buffer was taken before (lane 2) and after overnight incubation with detergents (lane 3). Centrifugation of the resulting suspension gives pellet 3 (lane 4), which contains insoluble POR, and supernatant 3 (lane 5) with POR incorporated in detergent micelles—the desired preparation. The volume of pellet 3 suspension was adjusted with water to match the volume of the supernatant 3 samples thus allowing for a direct comparison of the fractionation of POR between soluble and insoluble forms on the SDS-PAGE (lanes 4 and 5). The use of Triton X-100 (Figure 3.A) results in a small degree of POR extraction of 5-10% (ratio of the POR band intensity for lanes 5 and 4). This is despite the detergent concentration significantly exceeding its CMC value—the 2% v/v of Triton X-100 corresponds to 30 mM (compare to its CMC of less than 1 mM in Table 1).

**Table 1.** The CMC values of the evaluated detergents (based on reference data from Sigma-Aldrich and EMBL).

| Detergent | CMC (mM) |
| --- | --- |
| TritonX-100 | 0.2-0.9 |
| TritonX-114 | 0.2 |
| TritonX-405 | 69 |
| TWEEN20 | 0.06 |
| Brij35 | 0.09 |
| CHAPS | 6.4 |

**Figure 3.**
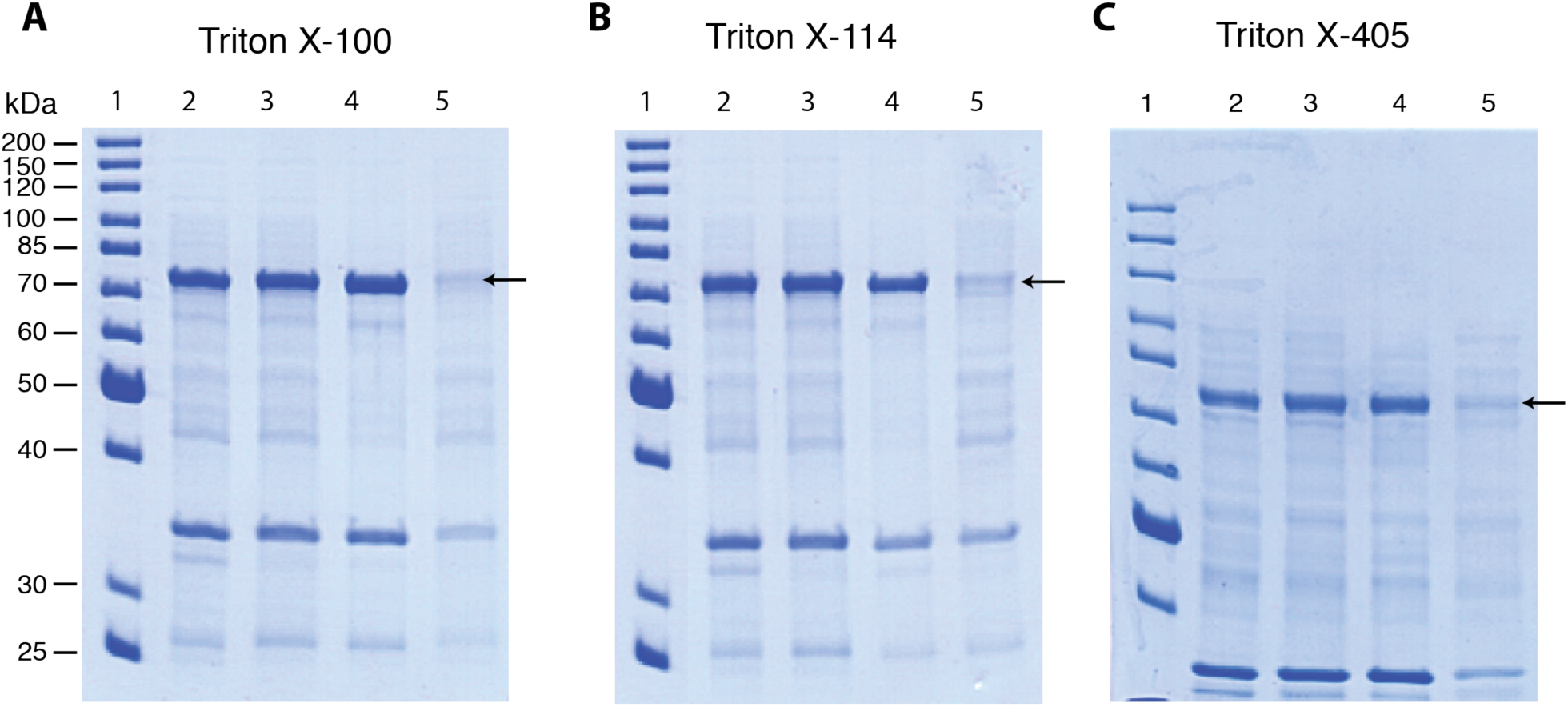
Solubilization of full-length POR by the Triton X-series detergents. In all panels, lane 1 is the PageRuler Unstained Protein Ladder; pellet 2 before and after overnight incubation with the detergent, lanes 2 and 3, respectively; pellet 3—insoluble POR fraction, lane 4; supernatant 3—solubilized POR, lane 5. The band of POR at 78 kDa is indicated by the arrow. All detergents were added to the solubilization buffers at 2% by volume.

Figure 3 panels B and C demonstrate the efficiency of POR extraction with different detergents of the Triton X series. The Triton X-series detergents are non-ionic molecules with a constant hydrophobic octylphenyl moiety and a variable polar polyoxyethanol region (Supporting Figure 1). The number of the polar ethylene oxide units in their hydrophilic tail (*n*) varies with the Triton X-114 having the smallest *n* of 7-8, the Triton X-100—intermediate *n* = 9-10, and with the Triton X-405 including the longest polyoxyethanol chain, *n* = 40. Correspondingly, hydrophobicity of a Triton detergent is increased with the smaller *n*: the Triton X-114 is more hydrophobic than the Triton X-100, while the Triton X-405 is the most hydrophilic of the three. Examination of Figure 3 reveals that POR extraction is relatively independent of the relative hydrophobicity of the Triton X detergents.

Our next extraction trial included detergents that had distinct molecular structures, TWEEN 20 and Brij 35. TWEEN 20 (Supporting Figure 2) is a polyoxyethylene sorbitan ester of a fatty acid. It is distinct from Triton in that its hydrophobic moiety is linear, as compared to the branched and aromatic octylphenyl of Tritons. The distinctive feature of Brij 35 (Supporting Figure 3) is an unbranched arrangement of a hydrophobic lauryl chain continuing into a hydrophilic polyoxyethylene chain. Figure 4 shows SDS-PAGE analysis of POR extraction from pellet 2 by solubilization buffers containing 2% TWEEN 20 or Brij 35. Comparison of relative distribution of POR between lanes 5 (soluble) and 4 (remaining insoluble) with the results for Triton X-100 in Figure 3.A indicates that the extraction yield is also relatively insensitive to the particular molecular structure of the non-ionic detergent.

**Figure 4.**
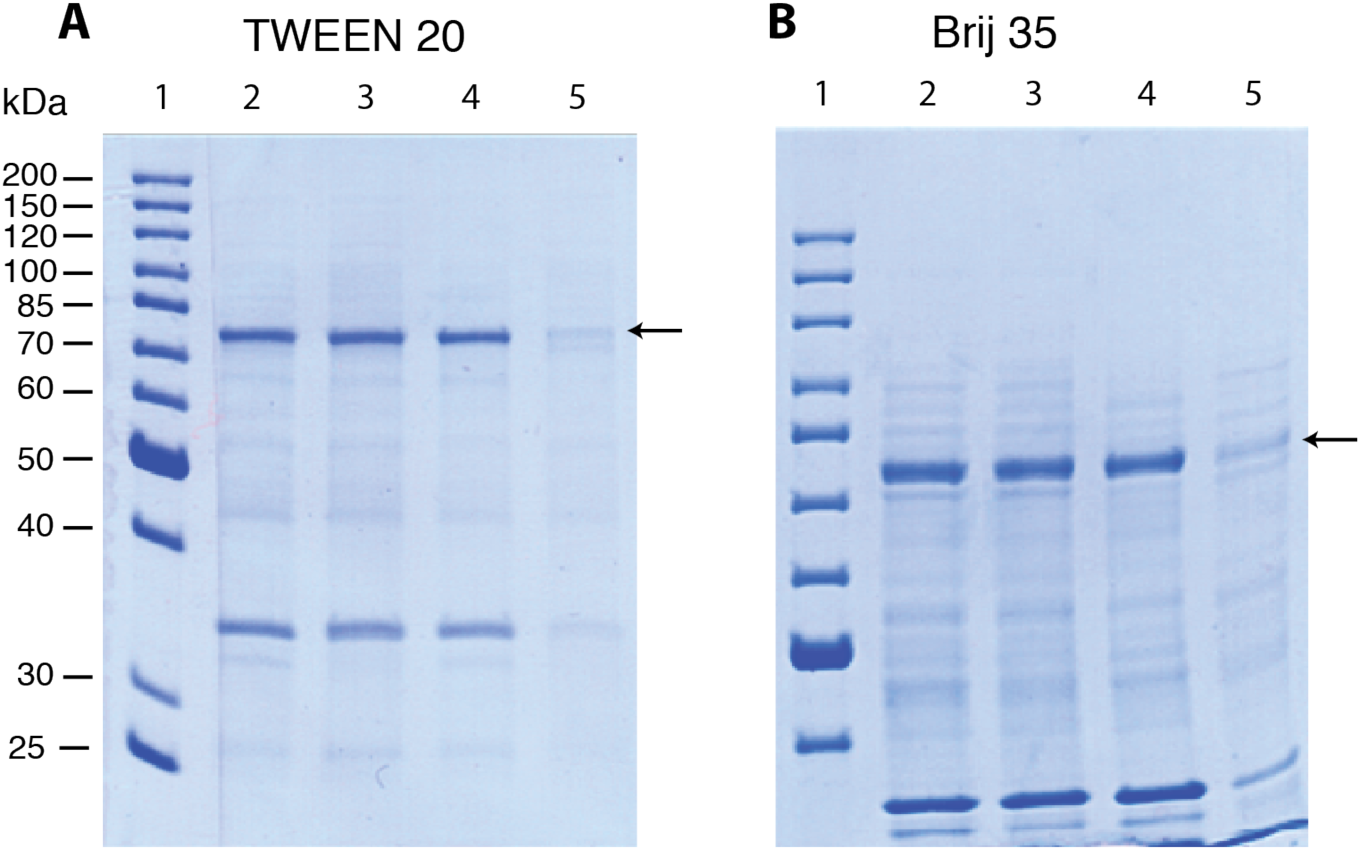
Solubilization of full-length POR by the TWEEN 20 and Brij 35. Lane assignments are the same as in Figure 3. The detergents were added to the solubilization buffers at 2% by volume.

The current literature indicates that a combination of detergents that have very distinct nature may result in an improved solubility of some membrane proteins. For example, Everberg et al. combined the zwitterionic detergent, Zwittergent 3-10, and the non-ionic detergent, TritonX-114, to solubilize mitochondrial membrane proteins obtained from the yeast *Saccharomyces cerevisiae.*^10^ The extent of extraction by this combination of detergents was very high and similar to that of SDS, which is known for its maximum extraction efficiency due to the highly charged nature of an SDS molecule. The Triton and Zwittergent are neutral detergents but in different ways: Triton X-114 has an uncharged head group, while the opposite charges of Zwittergent 3-10 cancel each other resulting in an overall uncharged molecule. Similar to Zwittergent, CHAPS and CHAPSO are zwitterionic detergents that have been widely used to solubilize membrane proteins.^11^

To test the effect of a detergent mixture on POR solubility, we combined CHAPS (Supporting Figure 4) with Triton X-100 to solubilize full-length POR. To reduce the overall solution viscosity, we used Triton X-100 at a lower concentration in this mixture (0.5%), therefore, we performed a separate reference extraction by 0.5% TritonX-100 alone as well. Figure 5 compares extraction of POR by Triton X-100 at 0.5% (v/v) alone and with CHAPS at 0.5% (w/v). Inspection of the POR distribution between lanes 4 and 5 of Figure 3.A and Figure 5. A reveals that reduction of Triton concentration from 2% to 0.5% leads to a small improvement in the extraction efficiency while addition of CHAPS, apparently, reduced POR solubility (Figure 5.B).

**Figure 5.**
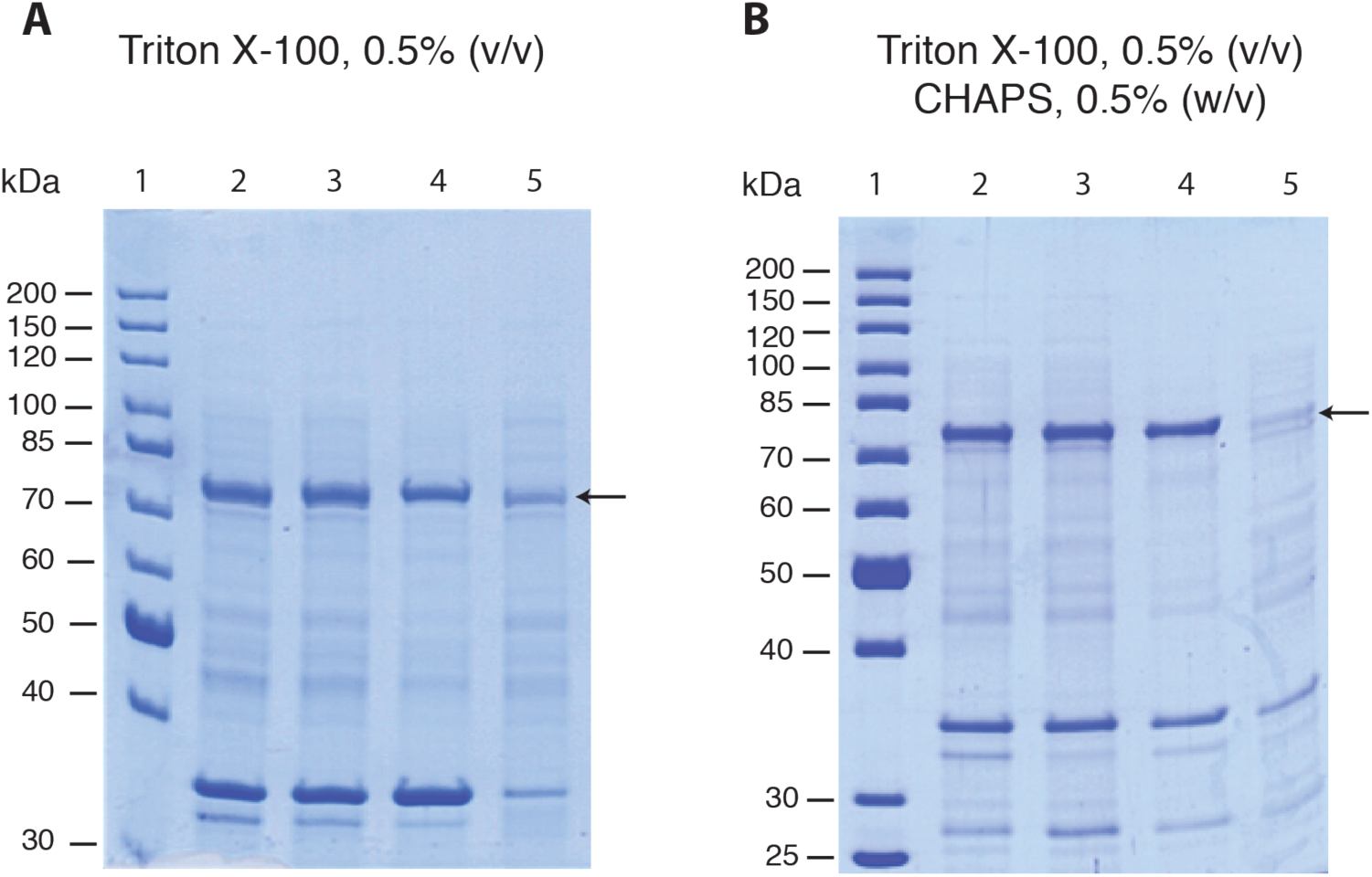
Solubilization of full-length POR by a mixture of detergents relative to the reference extraction with Triton X-100. Lane assignments are the same as in Figure 3

In the final chapter of this study, we evaluated several alkyl and polyamine additives that were reported to significantly improve extraction of some membrane proteins.^9^ Yasui et al. suggested that the cationic character of these additives allows them to interact with the negatively charged phosphates of phospholipids thus helping the nonionic detergent disrupt the phospholipid membranes and solubilize membrane proteins.^9^

In our trials shown in Figure 6, we complemented 0.5% Triton X-100 with either spermidine, spermine, putrescine, propylamine, or ethylamine hydrochlorides. All polyamines (panels B, C, and D) appeared to oppose solubilization. Propylamine (panel E) had little effect, while ethylamine (panel F) showed more protein in lane 5 than in the reference panel A. To ascertain the effect of ethylamine, we repeated the extraction in the presence of this additive and observed (Figure 7) that the overall extraction of POR is not improved relative to Triton X-100 alone. Careful inspection of Figure 6, panels A and F, reveals that ethylamine extraction gel in panel F had slightly greater overall sample load in all lanes. Therefore, the extraction efficiency estimated as a ratio of lane 5 to lane 4 is not really improved in panel F, which is in accord with the repeated experiment shown in Figure 7.

**Figure 6.**
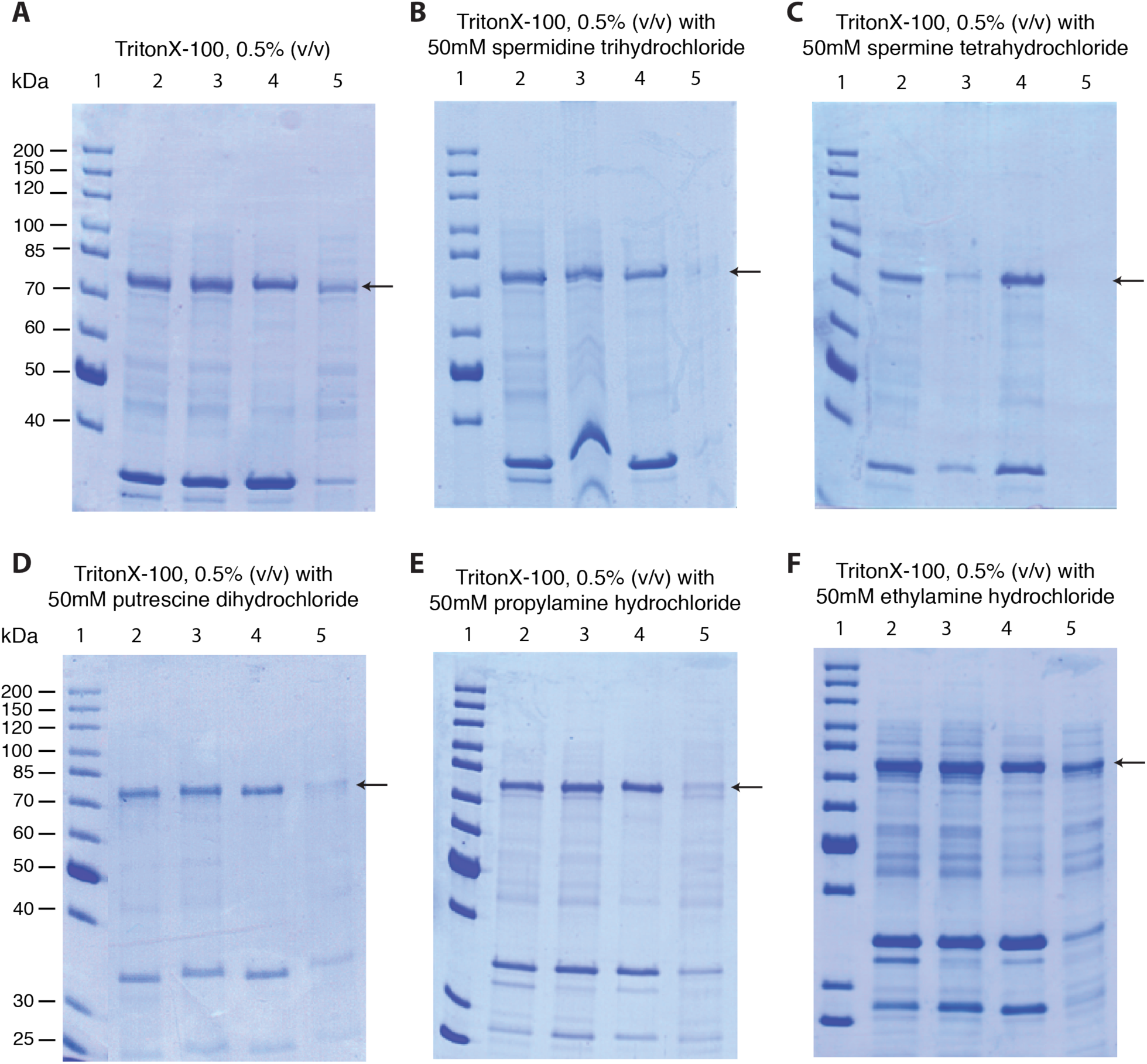
Solubilization of full-length POR by 0.5% Triton X-100 in the presence of alkyl and polyamines. Panel A is a reference experiment with 0.5% Triton X-100 alone (from Figure 5.A). Lane assignments are the same as in Figure 3.

**Figure 7.**
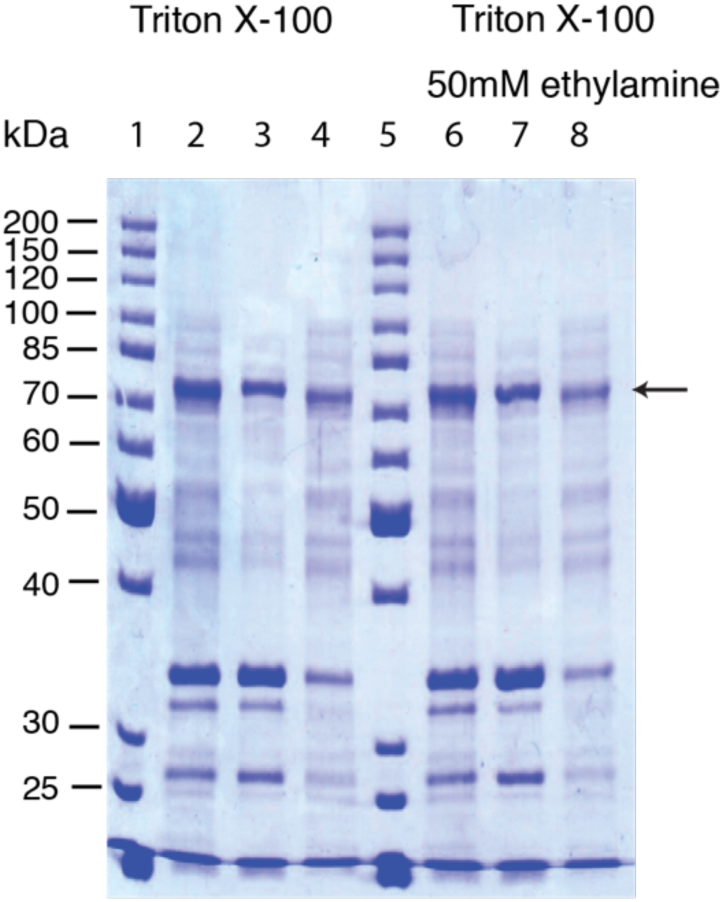
Solubilization of full-length POR by 0.5% Triton X-100 in the absence (lanes 2-4) and the presence of ethylamine (lanes 6-8). Resuspended pellet 2, lanes 2 and 6; insoluble fraction, lanes 3 and 7; solubilized POR, lanes 4 and 8; PageRuler Unstained Protein Ladder, lanes 1 and 5.

One possibility for the POR to remain insoluble in a detergent is if the protein forms large cross-linked aggregates. Our construct lacks native cysteines but includes two mutations that introduced solvent-exposed cysteine residues at the positions 157 and 517 (to allow for labeling by thiol-reactive probes). To prevent disulfide cross-linking, the solubilization buffer in this work included TCEP (the choice dictated by the use of Ni-chelation chromatography in the POR purification protocol^8^). However, TCEP is known to be relatively unstable in the phosphate buffers^12^, thus might not have provided a sufficient reducing capacity upon overnight incubations during the POR extraction step. Therefore, we performed an additional extraction experiment with Triton X-100 substituting 1 mM TCEP in the solubilization buffer with 1 mM DTT, which is stable in the presence of phosphates. Figure 8 shows that the replacement of TCEP for DTT had no effect on the extraction efficiency, which rules out the insufficient reduction of disulfide cross-linkages as a reason for insolubility of POR in the pellet 3.

**Figure 8.**
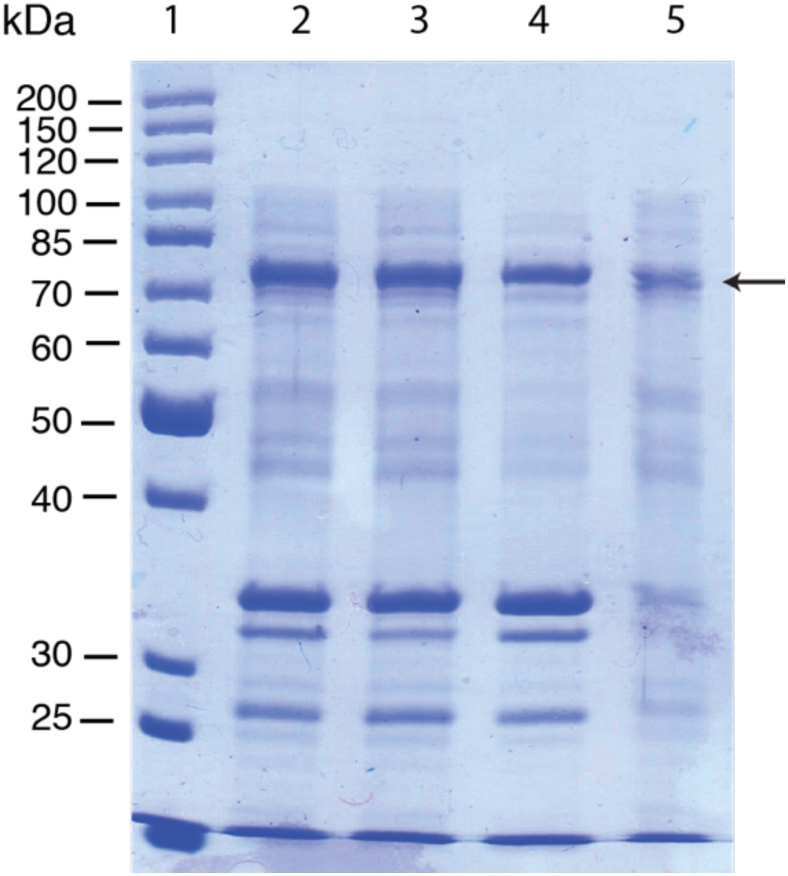
Solubilization of full-length POR by 0.5% Triton X-100 using DTT instead of TCEP as a reducing agent. Lane assignments are the same as in Figure 3.

In summary, we could not improve extraction of POR by broad variation of the composition or reducing capacity of the solubilization buffer. Therefore, we must conclude that in a typical extraction, we solubilize *all available* POR that is monomeric and properly anchored at the membrane. The remaining POR must be expressed in some unusual form, which is not the inclusion bodies (because they are removed with pellet 1). This POR fraction survives slow centrifugation but is pelleted by the fast speed (along with membrane vesicles) and remains insoluble in detergents. We hypothesize that this is an unfolded POR not associated with membranes (hence insensitive to detergents) and aggregated to a moderate extent. If this hypothesis is true, to improve the soluble POR yield, one needs to further optimize expression conditions to produce more of the properly folded monomeric protein. The key to success will be to gauge optimality of the expression conditions not by the *total quantity* of the overexpressed POR (like Figure 2) but by a *distribution* of POR between soluble and insoluble fractions in the detergent extraction step (supernatant 3 and pellet 3 in Figure 1).

## ACKNOWLEDGMENT

This work was supported by the National Institute of General Medical Sciences of the National Institutes of Health under award numbers R15 GM126528-01 to ELK; ARG acknowledges Eisch Research Fellowship during the academic year 2016-2017.

## Supporting Information

Supporting Figures and Tables.

## Supporting Information

**Supporting Figure 1.**
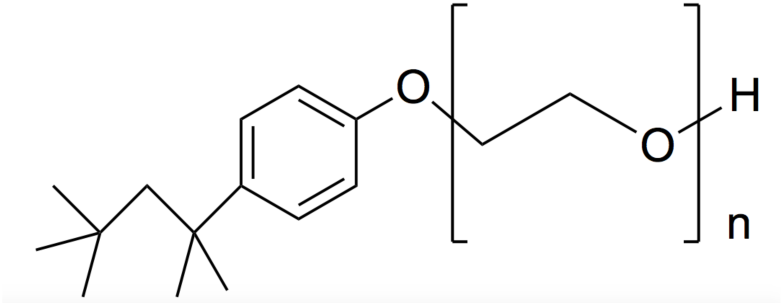
The general structure of the Triton X-series detergents. These detergents have the same hydrophobic moiety and differ in the number of the ethylene oxide units (n) in their hydrophilic tail.^14^

**Supporting Figure 2.**
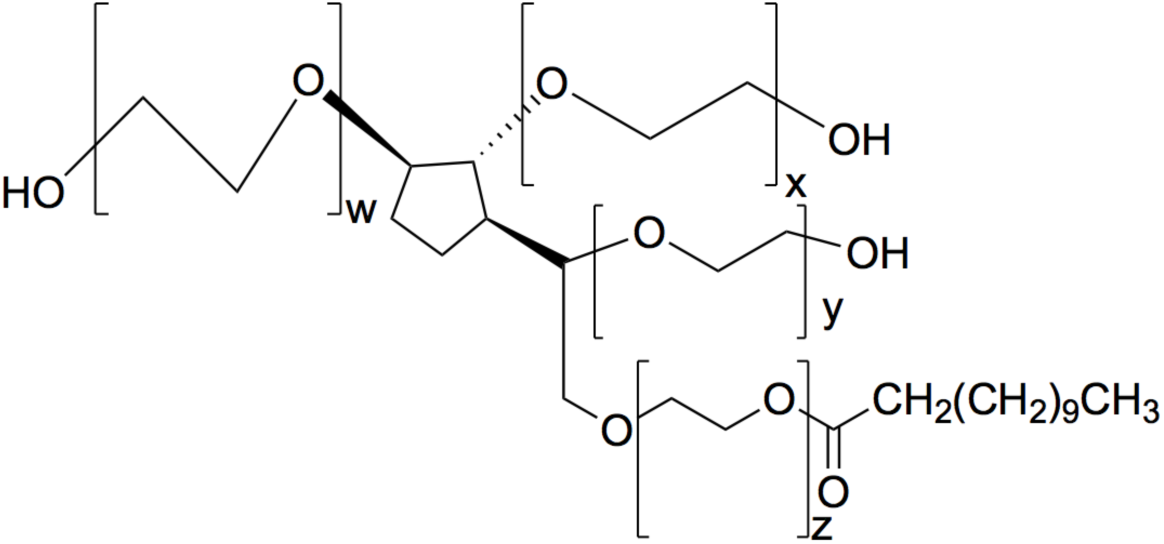
The structure of TWEEN 20. This non-ionic detergent has a total of 20 ethylene oxide units (x + y + w + z) and the lauric acid as the fatty acid tail.^15^

**Supporting Figure 3.**
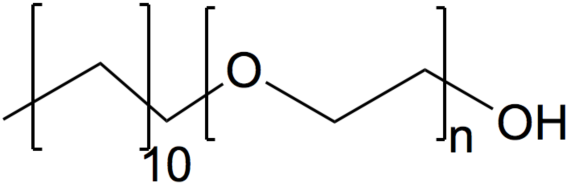
The structure of Brij 35. This non-ionic detergent has 23 ethylene oxide units (the “n” value) and has lauryl alcohol in its lipophilic portion.^16^

**Supporting Figure 4.**
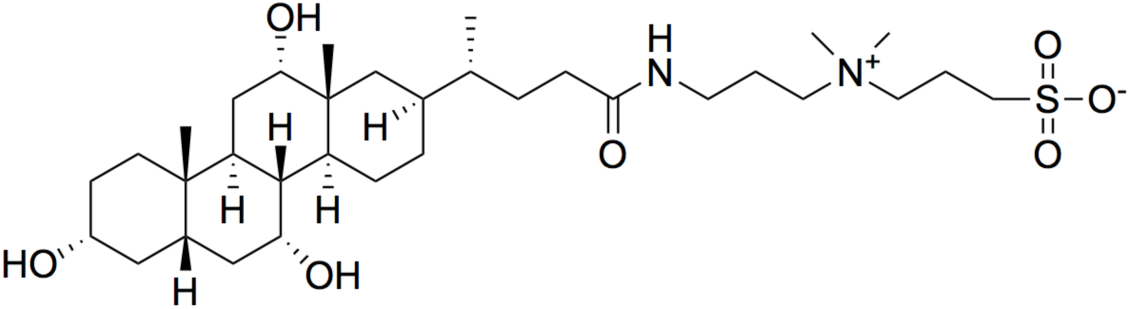
The structure of CHAPS. This zwitterionic detergent includes a negatively charged sulfonate in its hydrophilic region and an uncharged hydrophobic/hydrophilic cholic group.^17^

**Supporting Figure 5.**
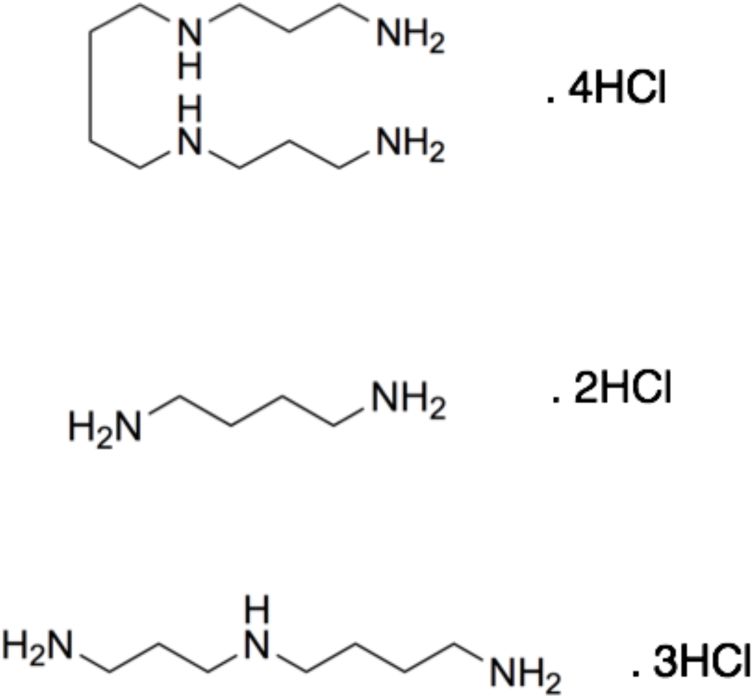
The structures of spermine tetrahydrochloride (top), putrescine dihydrochloride (middle), and spermidine trihydrochloride (bottom).^9^

